# Distinct Longitudinal Antibody Responses Following Primary Influenza Infection or Vaccination in Infants: A Prospective Birth Cohort Study

**DOI:** 10.64898/2026.09.28.755012

**Authors:** Devyani Joshi, Sanjeev Kumar, Veronika I. Zarnitsyna, Allison R. Burrell, Chelsea Rolhfs, Shamika Danzy, Tysheena Charles, Faith A. Mbadugha, Lingling Xu, Daniel C. Payne, Jacob E. Kohlmeier, Anice C. Lowen, Mary Allen Staat, Jens Wrammert

**Author notes:** Department of Pharmaceutical Sciences, School of Pharmacy, MCPHS University, Boston, MA, USA.

## Abstract

The longitudinal effects of initial influenza exposure on the magnitude, persistence, and antigen specificity of antibody responses during infancy remain incompletely understood. We investigated humoral antibody responses following primary influenza infection or seasonal vaccination in a prospective US birth cohort with weekly respiratory surveillance for influenza infection. Influenza-specific serum IgG binding to H1 and H3 haemagglutinin (HA) antigens, HA inhibition (HAI), and live-virus neutralizing antibody responses were assessed longitudinally. To minimize the influence of maternally derived antibodies, analyses of primary exposure responses were restricted to infants for whom influenza-specific maternal antibodies had declined below the assay detection limit before exposure. Antibody responses were compared following primary infection and vaccination and examined after subsequent annual vaccination. We found that maternal influenza-specific IgG declined rapidly during early infancy. Among infants without detectable pre-exposure influenza-specific antibodies, primary influenza infection was associated with greater post-exposure antibody responses and less measurable decline over follow-up than primary vaccination. These groups differed in age, calendar season, and exposure characteristics, limiting direct attribution of these differences to the route of exposure. In infants with previous influenza infection followed by vaccination, responses after subsequent vaccination initially showed greater binding to the previously infecting subtype among the tested HA antigens. With repeated vaccination, responses became more distributed across the tested antigens. Our findings reveal differences in the magnitude, persistence, and antigen specificity of antibody responses according to the nature and sequence of influenza exposure. By prospectively capturing early-life exposures, this study provides longitudinal evidence linking exposure history to subsequent humoral antibody responses.

## INTRODUCTION

The first encounter with influenza during childhood can influence the specificity and magnitude of antibody responses to subsequent influenza exposures. However, the longitudinal development of humoral immunity following primary influenza infection versus vaccination during infancy remains poorly defined^1^. Importantly, influenza remains a leading cause of respiratory illness and hospitalization in pediatric populations across the globe^2^. The virus’s ability to evade the immune response stems in part from annual antigenic drift, presenting an ongoing challenge to public health and vaccine development^3^. A longstanding obstacle in combating influenza is the complex and sometimes ambiguous nature of the acquired immune response. The historical term original antigenic sin (OAS) describes one formulation of this phenomenon, whereas contemporary studies use broader concepts including immune imprinting, back-boosting, and antigenic seniority^4^. OAS, which refers to the observation that an individual’s first encounter with an influenza virus shapes their humoral immune responses to all subsequent infections and vaccinations involving antigenically related viruses^4,5^. This initial exposure-associated antibody response creates a durable bias in immunological memory, in which memory B cell populations established in childhood dominate immune responses upon re-exposure to influenza, often at the expense of generating robust responses to novel epitopes encountered later in life ^6,7^. With each subsequent exposure, immune memory to strains encountered early in life is preferentially expanded, reinforcing a hierarchical antibody landscape in which back-boosted antibody responses prevail^5^. Although such memory can confer potent protection against closely related subtypes or clades, it may come at the cost of suboptimal immunity against circulating variants that escape neutralization due to antigenic drift. Thus, initial antigenic exposure is a double-edged sword: it can confer robust, long-lasting immunity against certain strains while limiting adaptive flexibility in the face of viral evolution^8,9^.

Studying the effects of initial antigenic exposure in adults is inherently challenging, given the complex history of multiple influenza exposures and repeated seasonal vaccinations, which obscure the immunological impact of the primary exposure event^10,11^. Following infants prospectively from birth provides an opportunity to characterize antibody responses associated with defined early-life influenza exposures before extensive exposure histories accumulate.

In this study, we leverage the Influenza IMPRINT (Immunological Memory to Prior INfluenza over Time), a prospective, longitudinal US birth cohort of mother-child pairs enrolled at Cincinnati Children’s Hospital Medical Center, to systematically characterize the associations between initial and sequential influenza exposures and humoral responses. Mothers were enrolled during their third trimester of pregnancy, and infants were followed from birth to at least three years of age. Influenza vaccination status was verified using state immunization registries and clinical provider records. Influenza infection history was obtained via weekly nasal swab samples collected by the parents, regardless of medical attention or symptom status. Longitudinal blood samples were obtained postnatally from cord blood, at week 6 of life, each summer, and acutely after influenza vaccinations or infections.

Here, we compared longitudinal antibody responses following primary influenza infection or primary vaccination and subsequently examined responses after repeated vaccination and after infection followed by vaccination. We further evaluate antibody responses to annual seasonal influenza vaccination in the absence of prior infection and characterize hybrid immunity in those who initially encountered influenza H1- or H3-infection and subsequently received annual seasonal influenza vaccination, in the same or subsequent influenza season. Our findings reveal differences in the magnitude, persistence, and antigen specificity of antibody responses according to the nature and sequence of influenza exposure. By prospectively capturing early-life exposures, this study provides longitudinal evidence linking exposure history to subsequent serum antibody trajectories.

## RESULTS

### Study cohort

Our samples included 91 US infants and children in the IMPRINT birth cohort who resided in the greater Cincinnati area, which includes Southwest Ohio and Northern Kentucky, and were intensively followed by researchers at Cincinnati Children’s Hospital Medical Center (CCHMC). Weekly parent-administered nasal swabs were collected regardless of symptoms to identify either symptomatic or asymptomatic influenza infections, and tested in real-time with multiplex reverse transcriptase polymerase chain reaction (RT-PCR) using the Flu SC2 assay (Centers for Disease Control and Prevention, Atlanta, Georgia)^12,13^ and NxTag Respiratory Pathogen Panel (RPP) (Luminex Molecular Diagnostics, Toronto, Canada)^14^. Nasal swabs positive for influenza by either the Flu SC2 assay or RPP were further tested with RT-PCR for Influenza A subtype and Influenza B lineage. Vaccinations for both mothers and their children were verified by the state immunization registry and/or clinical provider records. Longitudinal blood samples were collected postnatally from cord blood, at week 6 of life, each summer, and acutely after influenza vaccinations or infections.

The infants were stratified according to both their initial and subsequent influenza exposures. The overall study design is depicted in **Figure 1**. To evaluate antibody responses following primary influenza exposure, we selected infants with maternal binding antibody titers below the detection limit for all influenza antigens. This ensured that the maternal antibody titers did not interfere with the influenza exposure analysis. The groups included 17 infants with an initial H1-influenza infection and 22 infants with an initial H3-influenza infection. Here, we only included infants with at least 2 time points post-infection.

**Figure 1:**
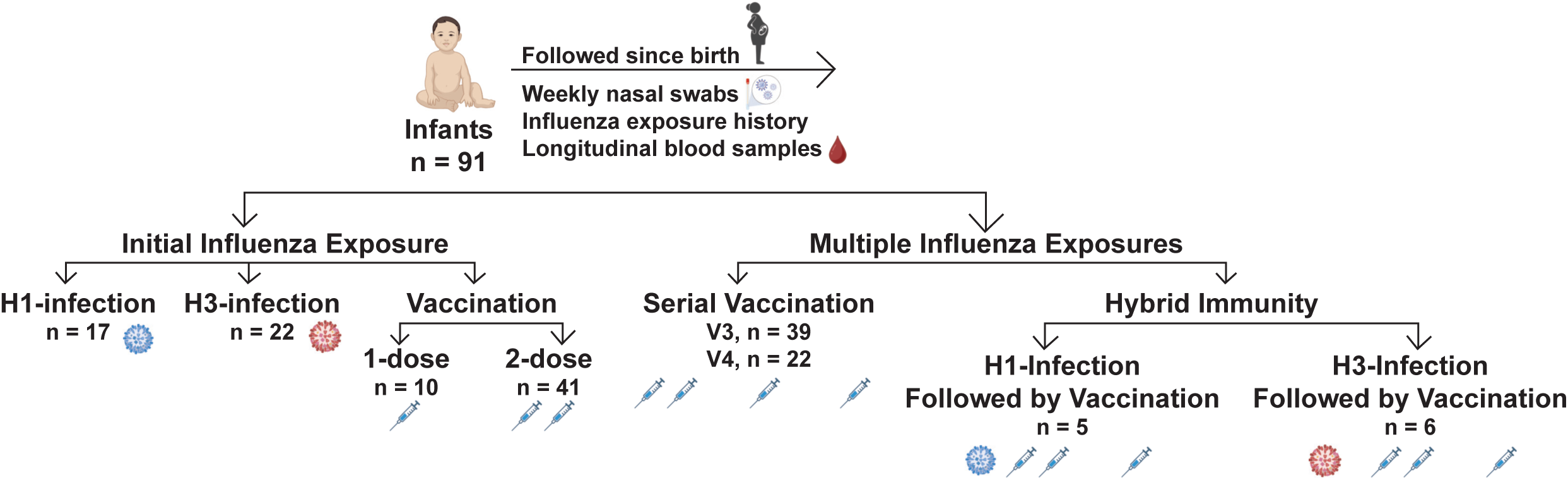
Study design. A total of 91 infants, followed since birth were included in the study. Influenza-specific binding and functional antibody responses were studied in these infants based on their influenza-exposure history, for the initial as well as multiple influenza exposures.

Additionally, infants who received the influenza vaccine as their first exposure were analyzed: 10 infants received only a single dose of the initial two-dose vaccine series, while 41 infants completed the full two-dose regimen during their first year (doses were administered one month apart). **Supplementary Figure 1** provides a comprehensive overview of these infants, detailing month and year of birth, timing of influenza vaccinations and infections, and all included longitudinal samples. Antibody responses following multiple influenza exposures were also investigated. This analysis included 39 infants who received three annual vaccine doses, and 22 infants who received four doses, with the initial two doses given a month apart in the first year of life. A subset of infants having “hybrid immunity” was also examined. This was defined as those who encountered a primary H1-(5 infants) or H3-(6 infants) influenza infection, followed by annual seasonal vaccination. **Supplementary Figure 2** outlines the full exposure and sampling history for these groups. The demographics and baseline characteristics of this cohort are described in **Table 1**.

**Table:** IMPRINT cohort demographics (n=91).

**Table: IMPRINT cohort demographics (n=91).**
| Variable | Values |  |
| --- | --- | --- |
| Gestational Age (Weeks) | 35-36 | 4 (4.4%) |
|  | 37 | 13 (14.3%) |
|  | 38-42 | 74 (81.3%) |
| Maternal Age at Enrollment (Years) | 18-24 | 8 (8.8%) |
|  | 25-34 | 58 (63.7%) |
|  | ≥ 35 | 25 (27.5%) |
| Gravidity | 1 | 21 (23.1%) |
|  | >1 | 70 (76.9%) |
| Fetal Sex | Male | 45 (49.5%) |
|  | Female | 46 (50.5%) |
| Pregravid BMI, kg/m<sup>2</sup> | Underweight | 3 (3.3%) |
|  | Healthy | 36 (39.6%) |
|  | Overweight | 19 (20.9%) |
|  | Obesity Class I | 17 (18.7%) |
|  | Obesity Class II | 7 (7.7%) |
|  | Obesity Class III | 9 (9.9%) |
| Delivery Mode | Vaginal | 56 (61.5%) |
|  | Operative Vaginal | 4 (4.4%) |
|  | Primary Cesarean | 15 (16.5%) |
|  | Repeat Cesarean | 16 (17.6%) |
| Maternal Race | White | 62 (68.1%) |
|  | Black | 22 (24.2%) |
|  | Asian | 2 (2.2%) |
|  | Other | 5 (5.5%) |
| Maternal Ethnicity | Hispanic | 5 (5.5%) |
|  | Non-Hispanic | 86 (94.5%) |
| Birthweight, kilograms | SGA | 5 (5.5%) |
|  | Not SGA | 86 (94.5%) |
| Maternal Influenza Vaccine Timing | First Trimester | 11 (12.1%) |
|  | Second Trimester | 16 (17.6%) |
|  | Third Trimester | 29 (31.9%) |
|  | No Vaccine During Pregnancy | 35 (38.5%) |
| Breastfeeding Duration | 0-5 Months | 36 (39.6%) |
|  | 6-11 Months | 23 (25.3%) |
|  | 12 Months or More | 32 (35.2%) |

### Maternal influenza-specific antibodies decline rapidly during early infancy

We examined the magnitude and persistence of maternal influenza-specific binding antibodies in infants at birth and up to 600 days post-birth who had no documented influenza exposure during this interval. A total of 58 infants born to influenza-vaccinated mothers in the influenza season 2020-2021 were included in this analysis (**Figure 2A**). We evaluated influenza-specific antibody binding titers at birth against four hemagglutinin (HA) antigens: H1/Hawaii/2019, H3/HongKong/2019, B/Phuket/2013, and B/Washington/2019, which corresponded to the vaccine strains for the 2020-2021 influenza season. All infants demonstrated robust influenza-specific antibody binding titers at birth, exceeding the limit of detection for all four antigens. There was no significant difference in the magnitude of these antibody titers at birth across trimesters in which mothers received influenza vaccination (**Figure 2B**), consistent with other studies indicating that the timing of immunization within pregnancy generally does not markedly affect infant antibody level^15,16^. The durability of maternally derived influenza-specific IgG responses was characterized by a rapid post-birth decline, with quantified half-lives of 34 (95 % confidence interval [95 % CI] [26, 51]), 35 (95 % CI [29, 43]), 40 (95 % CI [35, 45]), and 42 (95 % CI [35, 50]) days for binding antibodies against H1/Hawaii/2019, H3/Hong Kong/2019, B/Phuket/2013, and B/Washington/2019 HA antigens, respectively, as shown by an exponential decay model (**Figure 2C**). These findings demonstrate a rapid decline in maternally derived influenza-specific IgG during early infancy, defining the period over which passively acquired antibody levels decrease substantially. This observation underscores the importance of maintaining age-appropriate influenza immunization as maternal antibody levels decline.

**Figure 2:**
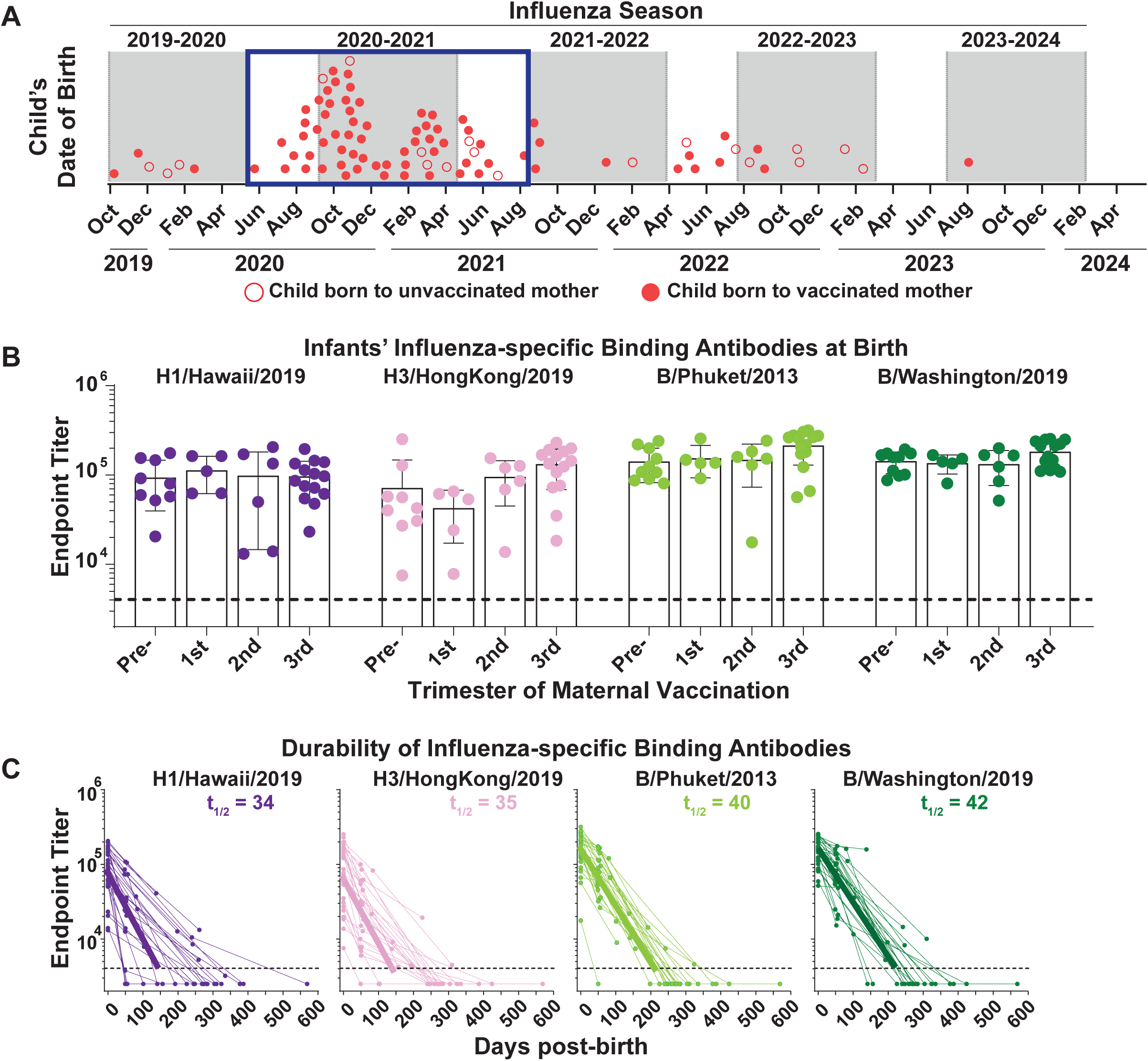
Influenza-specific maternal binding antibody responses in infants. (A) Child’s date of birth, the box represents the children included in the analysis for influenza-specific maternal binding antibody responses. (B) The influenza-specific binding antibody responses in the children at birth to H1/Hawaii/2019, H3/HongKong/2019, B/Phuket/2013, and B/Washington/2019 influenza HA antigens as determined by ELISA assay and expressed as Endpoint Titer. The children are categorized based on the receipt of maternal influenza vaccination trimester. (C) Durability of influenza-specific maternal binding antibody responses in infants to H1/Hawaii/2019, H3/HongKong/2019, B/Phuket/2013, and B/Washington/2019 influenza HA antigens as determined by ELISA assay and expressed as Endpoint Titer, along with their respective half-lives and decay curves. The dotted lines represent the limit of detection, defined as average + 3SD of the seronegative samples.

### Primary influenza infection and vaccination are associated with distinct longitudinal antibody responses in early life

We assessed both the magnitude and durability of influenza-specific binding and functional antibody responses in infants and children following their initial influenza exposure, whether through influenza infection with H1- or H3-strains or administration of the 1- or 2-dose influenza vaccine series. For this analysis, 10 infants received a single dose of the recommended two-dose influenza vaccine regimen in their first year, while 41 infants completed the full two-dose series, spaced approximately one month apart, during the 2021–2022 influenza season. In addition, infants who experienced their first exposure by H1- or H3-influenza infection during the 2021–2022 (9 infants: H3), 2022–2023 (13 infants: H3; 9 infants: H1), or 2023–2024 (6 infants: H1) seasons were included (**Figure 3A**). To measure the immune response, we quantified IgG-binding antibodies targeting H1/Wisconsin/2019, H1/Wisconsin/2022, H3/Tasmania/2020, and H3/Darwin/2021 hemagglutinin (HA) antigens, each corresponding to the specific strains relevant to the infant’s initial infection or vaccination season.

**Figure 3:**
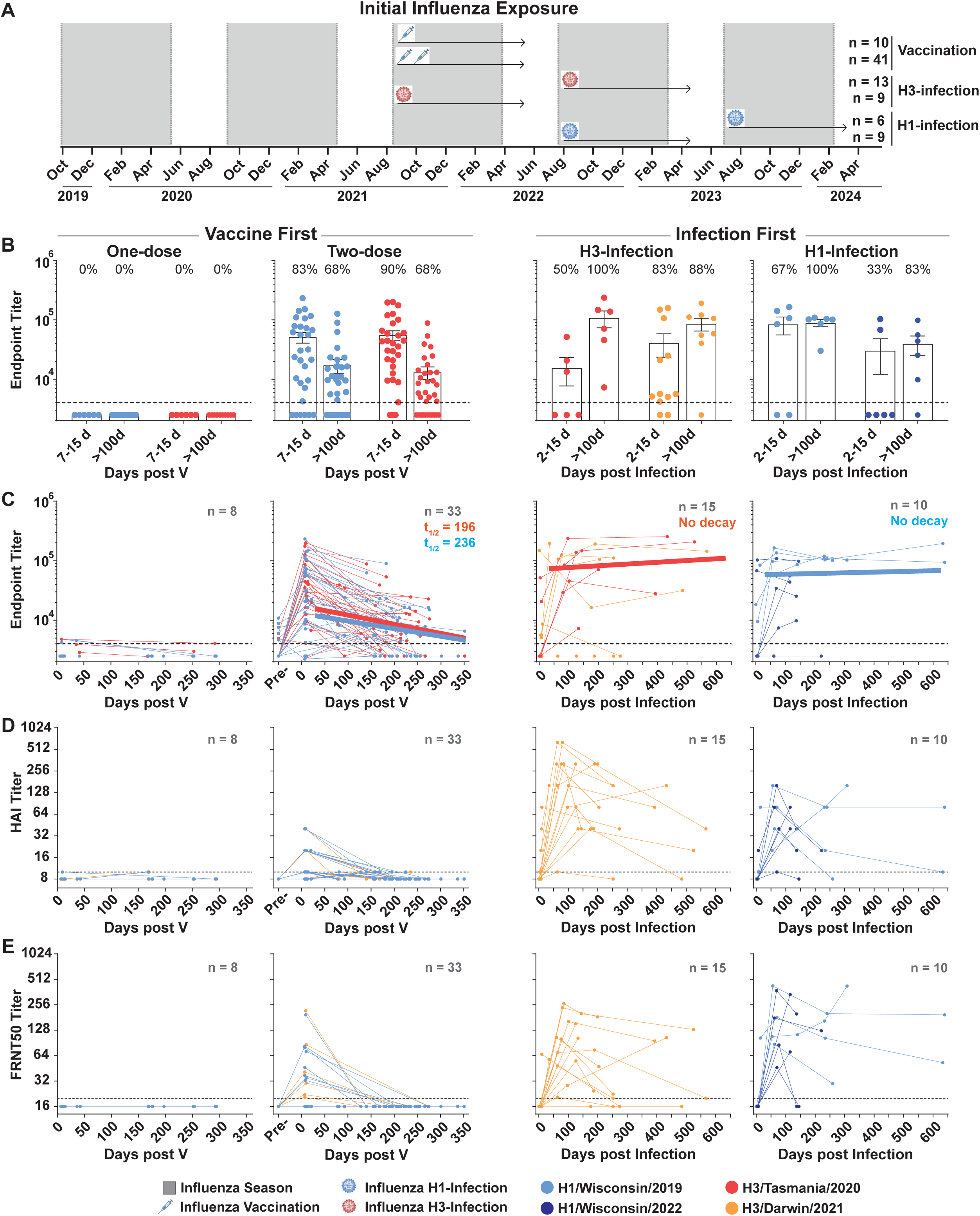
Binding and functional antibody responses to initial influenza exposures in infants. (A) Infants included in the analysis for initial influenza exposure, one-dose vaccination, two-dose vaccination (1 mo apart), H3-infection, or H1-infection. (B) Magnitude of influenza-specific binding antibody response at acute and convalescent time point post initial influenza exposure in infants as determined by ELISA assay and expressed as Endpoint Titer. (C) Durability of influenza-specific antibody responses in infants to H1, and H3 influenza HA antigens, along with their respective half-lives and decay curves. All observations are shown; exponential decay was fitted only to measurements obtained after day 28 from individuals with more than 2 post-day 28 measurements. (D) Durability of influenza-specific HAI titers in infants to H1, and H3 influenza HA antigens. (E) Durability of influenza-specific neutralizing antibody responses in infants to H1, and H3 influenza HA antigens.

In this cohort, a single dose of influenza vaccine as the initial exposure was not associated with detectable binding antibody responses to the tested H1 or H3 HA antigens at the sampled acute (7–15 days post-vaccination) or convalescent (>100 days post-vaccination) timepoints. In contrast, the two-dose vaccine schedule resulted in robust antibody responses, with 83% of infants mounting detectable responses against H1/Wisconsin/2019 HA, and 90% against H3/Tasmania/2020 HA antigens at the acute phase following the second dose. However, these vaccine-elicited responses declined rapidly, with only 68% of infants retaining detectable titers against both HA antigens at the convalescent timepoint (>100 days post-vaccination). Analysis of infants who experienced primary infection with H3 or H1 strains revealed that antibody responses were sometimes absent during the acute phase (2–15 days post-infection), likely because sampling occurred too early relative to seroconversion. Nevertheless, 50% and 83% of infants showed detectable titers against H3/Tasmania/2020 and H3/Darwin/2021 HA, respectively, after H3 infection, and 67% and 33% showed titers against H1/Wisconsin/2019 and H1/Wisconsin/2022 HA, respectively, after H1 infection. Unlike vaccine-induced responses, infection-induced antibody responses showed less measurable decline over the available follow-up, as evidenced by the convalescent data: 100% and 88% of infants retained detectable titers against H3/Tasmania/2020 and H3/Darwin/2021 HA, while 100% and 83% retained titers against H1/Wisconsin/2019 and H1/Wisconsin/2022 HA, respectively (**Figure 3B**).

We further characterized the durability of both binding and functional antibody responses - specifically HAI titer and FRNT50 titer - following initial influenza exposure in infants. Infants who received a single dose of influenza vaccine consistently exhibited antibody titers below the detection threshold at all post-vaccination timepoints. Thus, no measurable binding or functional antibody response was detected after a single vaccine dose in the sampled infants.

Among infants receiving the two-dose primary vaccine series, antibody concentrations declined over the follow-up period, with estimated half-lives of 236 days (95% CI, 130–1219) for H1 and 196 days (95% CI, 121–519) for H3, as calculated using an exponential decay model. Infants experiencing primary influenza infection with H1 or H3 strains exhibited markedly more durable binding and functional antibody responses specific to the infecting viral antigens. This durability was reflected in longer binding antibody durability after infection (no measurable decay against H3 and H1 HA antigens by exponential decay analysis) as compared to responses following vaccination. This pattern extended to functional immunity: both HAI and FRNT50 responses showed a smaller decline following infection than after primary vaccination over the available follow-up period, underscoring the pronounced difference in antibody longevity by exposure type (**Figure 3C, 3D, and 3E**).

Together, these findings indicate that primary influenza infection and primary vaccination were associated with different longitudinal trajectories of binding and functional antibody responses in this cohort. The observed differences should be interpreted in the context of differences in age, season, infecting strain, and other exposure characteristics between the groups.

### Repeated annual influenza vaccination is associated with increasing antibody responses in infants and young children

We further examined the immune response to repeated annual seasonal influenza vaccination in multiple exposures in infants and children. This analysis only included infants and children who received two doses of influenza vaccine (administered one month apart) in the 2021-2022 season (n=41), a third dose (V3) in 2022-2023 (n=39), and a fourth dose (V4) in 2023-2024 (n=22) (**Figure 4A**). We measured the magnitude of both binding and functional antibody responses, including hemagglutination inhibition (HAI) and focus reduction neutralization (FRNT50) titers at the acute timepoint (7–15 days post-vaccination) following each vaccination against the corresponding H1- and H3-HA antigens of each influenza season. Both binding and functional antibody responses against H1 and H3 HA antigens showed progressive increases in magnitude with an increasing number of annual vaccine doses. As previously described, after V2, 83% and 90% of infants registered binding antibody responses above the limit of detection against H1/Wisconsin/2019 and H3/Tasmania/2020 HA antigens, respectively. This proportion rose further with subsequent vaccinations: after V3, 96 % (H1) and 100 % (H3), and after V4, 100 % (H1) and 95 % (H3) of infants had detectable binding antibodies to the season-matched antigens at the acute timepoint. A similar upward trend was observed for functional antibody responses, HAI, and FRNT50 titers. After V2, 43 % (H1) and 40 % (H3) infants registered HAI titers, and 23 % (H1) and 30 % (H3) infants had neutralizing antibody titers above the limit of detection. Post V3, 100 % (H1) and 93 % (H3) infants showed HAI titers, and 85 % (H1) and 81 % (H3) infants showed detectable neutralizing antibody titers. Similar response rates were maintained after V4, with 94 % (H1) and 94 % (H3) infants showing HAI titers and 82 % (H1) and 82 % (H3) infants showing neutralization titers above the detection limit, supporting an association between repeated annual vaccination and increased antibody response magnitude in this cohort (**Figure 4B**). To assess the antibody durability, we evaluated binding and functional antibody titers up to 350 days post-vaccination. After V2, half-lives of antibodies against H1/Wisconsin/2019 and H3/Tasmania/2020 HA antigen were 236 (95 % CI [130, 1219]) and 196 (95 % CI [121, 519]) days, respectively. Whereas after V3, the antibody half-lives were 158 (95% CI [119, 235]) and 103 (95% CI [58, 457]) days against H1/Wisconsin/2019 and H3/Darwin/2021 HA antigens, respectively (**Figures 4C-D**). Here, we could not evaluate the durability of binding antibody responses post-V4 due to inadequate sampling points post-vaccination. Together, these data demonstrate that repeated annual influenza vaccination in infancy and childhood enhances the magnitude of both binding and functional antibody responses, underscoring the importance of ongoing annual vaccination for sustained antibody-mediated immunity against influenza.

**Figure 4:**
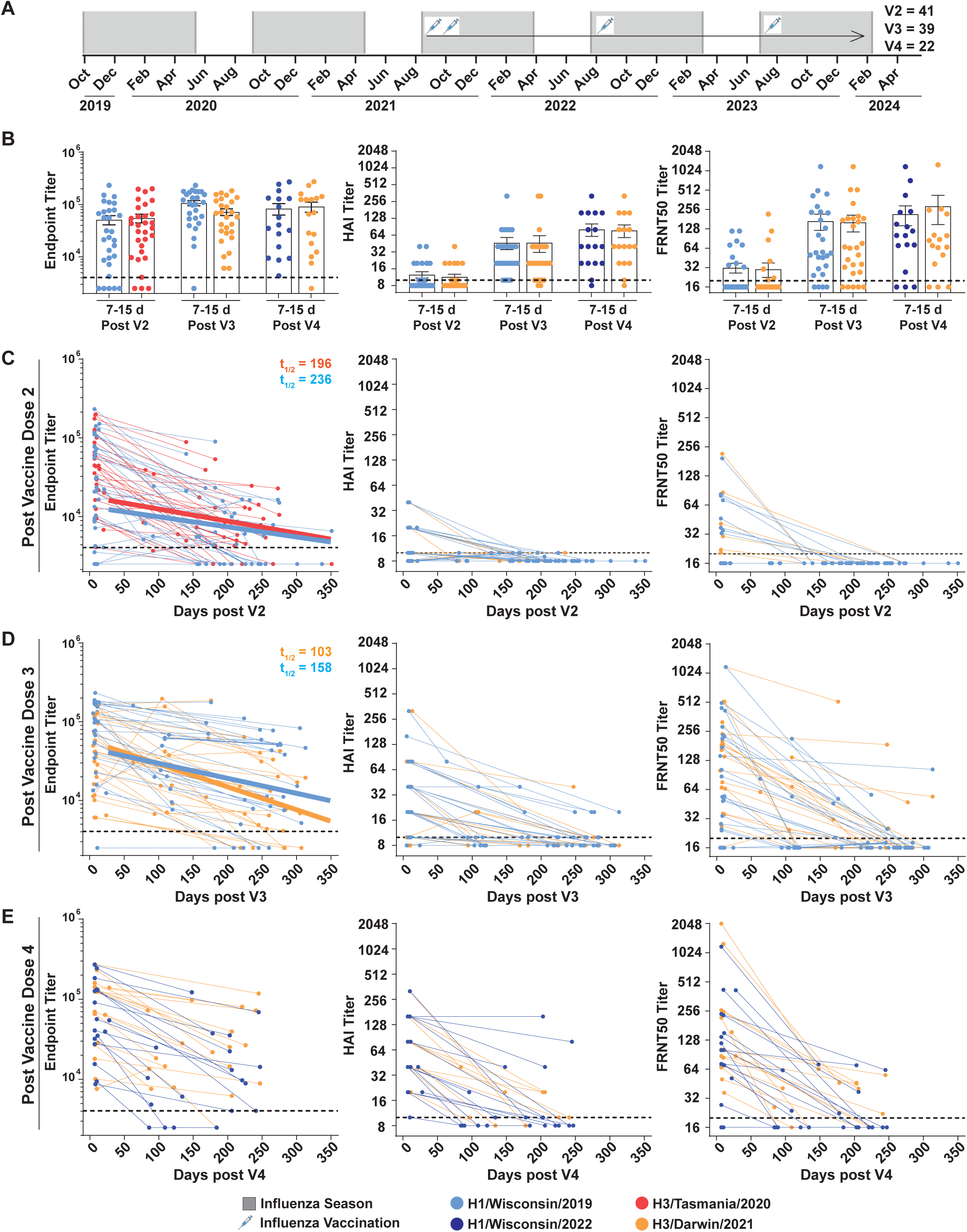
Binding and functional antibody responses to yearly seasonal influenza vaccination in infants. (A) Infants included in the analysis for initial influenza exposure as serial vaccination. This shows the number of infants receiving 2-dose (1 mo apart), 3 doses, and 4 doses of seasonal influenza vaccines. (B) Magnitude of influenza H1- and H3-specific binding, functional & neutralizing antibody responses in infants post 2, 3, and 4-doses of seasonal vaccines as determined by ELISA, HAI assay, and neutralization assay, respectively. (C) Durability of influenza-specific antibody responses in infants to H1, and H3 influenza HA antigens, along with their respective half-lives and decay curves post 2-doses of seasonal influenza vaccination. (D) Durability of influenza-specific antibody responses in infants to H1, and H3 influenza HA antigens, along with their respective half-lives and decay curves post 3-doses of seasonal influenza vaccination. (E) Durability of influenza-specific antibody responses in infants to H1, and H3 influenza HA antigens, post 4-doses of seasonal influenza vaccination.

### Prior influenza infection is associated with initially exposed strain-biased antibody responses following subsequent vaccination

We investigated hybrid immune responses in infants who, after experiencing prior influenza H3 (n=6) or H1 (n=5) infection, subsequently received annual seasonal influenza vaccinations (**Supplementary Figure 2B**). Here, we included all infants with influenza H1- or H3-infection as a first exposure followed by seasonal vaccination. Only a subset of these infants was included in the infection-first group of infants analyzed in **Figure 3**. **Figure 5A** displays longitudinal binding antibody titers against H1/Wisconsin/2019, H3/Darwin/2021, B/Phuket/2013, and B/Washington/2019 HA antigens in three representative donors: CC0282, who received only annual influenza vaccinations; CC0708, who had an initial H3 infection followed by annual influenza vaccinations; and CC0929, who had an initial H1 infection followed by annual influenza vaccinations. In infants who received repeated vaccinations in the absence of infection, binding antibody responses of comparable magnitude were generated against all four HA antigens. Pie charts in **Figure 5A** illustrate a roughly equal distribution of these binding antibodies at the acute time points V2 and V3. Conversely, infants with initial H3 infection exhibited antibody responses post-first vaccine dose that were heavily skewed toward the infecting antigen, with approximately 88% of the summed binding signal across the four tested HA antigens directed toward H3 in this representative infant. Similarly, infants initially infected with H1 generated responses after the first vaccination that were predominantly against the H1 antigen, comprising about 87% of the summed binding signal across the four tested HA antigens, with this response directed toward H1 in this representative infant. This antigenic skew persisted but gradually normalized with further annual vaccinations, although a higher proportion of antibodies remained directed toward the initial infecting strain through the third vaccine dose. Specifically, H3-infected and vaccinated infants had 52% and 38% of binding antibodies targeting H3 after the second and third vaccine doses, respectively, whereas H1-infected infants showed 60% and 39% of binding antibodies targeting H1 at the same time points. We then compared the magnitude of binding and functional antibody responses across first, second, and third vaccine doses in infants with prior H3 or H1 infection (**Figure 5B**). These infants mounted strong binding and functional antibody responses against their infecting strain post-V1, which persisted through subsequent doses. Antibody responses against the alternate strain (H1 for prior H3-infected and H3 for prior H1-infected infants) were initially negligible after the first vaccination but increased gradually with subsequent vaccine doses. By V3, these responses were comparable in magnitude to those of infants receiving multiple vaccinations without prior infection, indicating that vaccine-induced immunity against non-infecting strains develops robustly regardless of infection history. Together, these data underscore the importance of annual seasonal influenza vaccination in early life, irrespective of prior infection status, to achieve broad and durable protection against circulating influenza virus strains.

**Figure 5:**
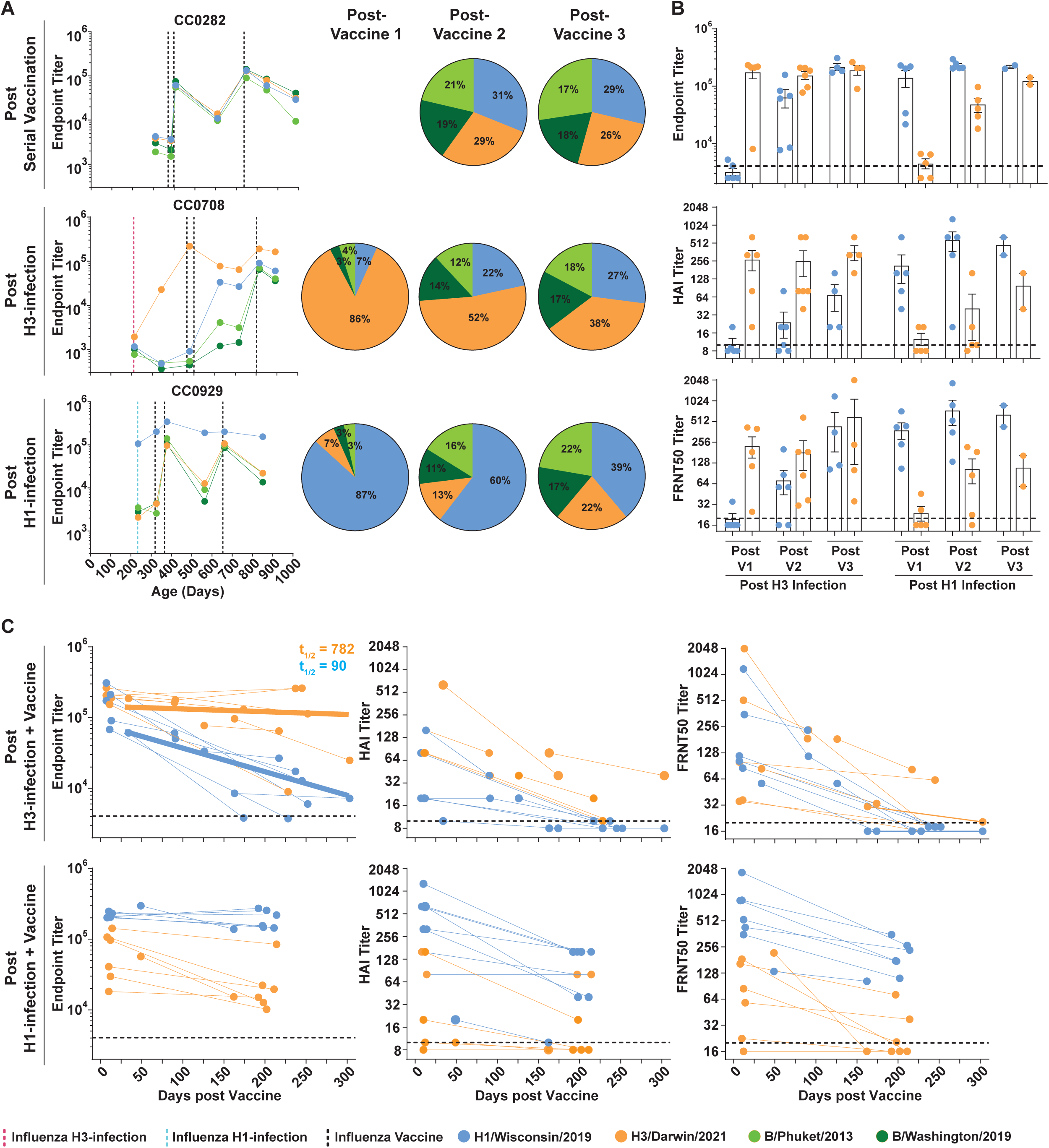
Binding and functional antibody responses in infants with influenza infection followed by serial vaccination. (A) Representative infant donors having influenza exposure as serial vaccination, influenza H3-infection followed by serial vaccination, and influenza H1-infection followed by serial vaccination, along with the antibody distribution against 4 different influenza antigens post 2nd, and 3rd dose of vaccine. (B) Magnitude of influenza-specific binding antibody response as determined by ELISA assay and expressed as Endpoint Titer, HAI Titer, and Neutralizing antibody titer post serial vaccination in infants with prior influenza infection. (C) Durability of influenza-specific binding antibody responses, HAI titer, and neutralizing antibody titer in infants post serial vaccination with prior influenza infection to H1, and H3 influenza HA antigens, along with their respective half-lives and decay curves.

### Prior infection is associated with greater persistence of post-vaccination antibody responses to the previously encountered subtype

We next assessed the durability of binding and functional antibody responses following vaccination in infants with prior influenza infection with H3 (n=6) or H1 (n=5) (**Figure 5C**). Prior infection was associated with greater durability of post-vaccination antibody responses against the homologous (infecting) strain than against the heterologous strain. Specifically, vaccination in infants with prior H3 infection yielded a binding antibody half-life of 782 (95 % CI [156, inf]) days against the H3 antigen, significantly longer than 90 (95 % CI [60, 178]) days against the H1 antigen, as estimated using an exponential decay model. A similar pattern was observed for functional antibody responses, with more durable responses elicited against the previously encountered strain than against the non-infecting strain. Taken together, these data suggest that early-life infection was associated with greater persistence of post-vaccination antibody responses against the previously encountered antigen than against the heterologous antigen. This pattern is consistent with an exposure-history-associated bias in subsequent antibody responses.

## DISCUSSION

The longitudinal design of this birth cohort allowed us to characterize serum antibody responses from the period of maternally derived immunity through primary influenza exposure and subsequent vaccination. Weekly respiratory surveillance and serial blood collection provided a defined record of influenza infections and vaccinations and enabled assessment of antibody responses before and after exposure. This design addresses an important limitation of studies in older children and adults, in whom early exposure histories are often difficult to reconstruct accurately. Previous cross-sectional studies in infants have identified differences in influenza antibody responses by exposure history ^18–21^, whereas longitudinal studies beginning before substantial influenza exposure remain relatively limited ^22,23^.

We first observed a rapid decline in maternally derived influenza-specific antibodies during early infancy, consistent with previous studies ^29,34,35^. The estimated half-lives across the influenza antigens tested were approximately 34-42 days, indicating that maternally derived serum antibody levels decline substantially during the first months of life. These findings define the period over which maternal antibody wanes but do not establish the duration or clinical effectiveness of passive protection. Breastfeeding history was not incorporated into the present analysis, and therefore, potential antibody exposure through breastmilk could not be distinguished from transplacentally acquired antibody.

Among infants whose influenza-specific antibodies were below the assay detection limit before their first documented exposure, primary influenza infection was associated with greater-magnitude binding and functional antibody responses and less measurable decline over follow-up than primary vaccination. Following the two-dose primary vaccination regimen, antibody responses increased at the early post-vaccination timepoint but subsequently declined, with estimated binding-antibody half-lives of 236 days for H1 and 196 days for H3. In contrast, no measurable decay was detected in the corresponding antibody responses following primary H1 or H3 infection over the available follow-up period. The infection-associated responses also showed greater persistence in HAI and FRNT50 measurements. These observations indicate that the two exposure groups had different longitudinal serum antibody trajectories in this cohort.

Infection and vaccination were not randomly assigned, and the groups differed in age at exposure, calendar season, and exposure characteristics. Differences in immune maturation, circulating or vaccine strains, and other participant characteristics could therefore contribute to the observed differences in antibody magnitude or persistence. In addition, the infecting virus was classified at the subtype level, and this does not establish antigenic identity between the infecting virus and each HA antigen used in the serologic assays. Some post-infection samples were obtained only 2-15 days after infection and may have been collected before seroconversion, further complicating comparisons of early responses. Thus, the observed differences should not be interpreted as demonstrating that infection intrinsically produces superior immunity to vaccination.

Repeated vaccination was associated with progressively greater binding and functional antibody responses across successive vaccine exposures. Among children who received the two-dose primary series followed by annual vaccination, responses to the season-matched H1 and H3 antigens increased from V2 to V3 and remained high after V4. However, V3 and V4 were nested within the longitudinal cohort, included fewer participants than V2, and had vaccine antigens that differed across seasons. Thus, the progressive increase cannot be attributed solely to the cumulative number of vaccine doses, because age, calendar season, and antigenic differences also changed over time. Nevertheless, the longitudinal pattern demonstrates that repeated vaccination was associated with increasing serum antibody responses in these children.

The analysis of children with influenza infection followed by vaccination provides additional evidence that exposure sequence is associated with subsequent antibody specificity. In the small subgroup of 11 children, including six with prior H3 infection and five with prior H1 infection, post-vaccination binding responses were initially enriched for the previously encountered subtype among the four HA antigens tested. With additional vaccination, this distribution became less dominated by the previously infecting subtype and more broadly distributed across the HA antigens tested. Because **Figure 5A** presents three representative participants and the analysis included only 11 children, these findings should be considered exploratory. Importantly, the assay measured serum binding to a defined set of HA antigens rather than the complete antibody or B-cell repertoire. Thus, the findings support an exposure history-associated pattern in measured antigen reactivity but do not establish a broader change in the underlying antibody repertoire.

Prior infection was also associated with greater persistence of post-vaccination antibody responses against the previously encountered subtype in this small subgroup. For example, following prior H3 infection, the estimated H3 binding-antibody half-life was 782 days, compared with 90 days for H1; however, the confidence interval for the H3 estimate was wide, extending to infinity. This estimate should therefore be interpreted as evidence of limited measurable decay over the available observation period, rather than of indefinite antibody persistence. Similar subtype-associated patterns were observed for functional antibody responses. The cellular basis of these differences cannot be determined from the present study. Previous studies have implicated memory B cells and long-lived plasma cells in durable influenza antibody responses24-27, but these populations, as well as antibody affinity, clonal lineages, germinal-center responses, and T-cell responses, were not measured here.

We did not have sufficient power to determine whether responses to heterologous HA antigens differed between children whose first exposure was infection followed by vaccination and those whose first exposure was vaccination. The absence of a statistically significant difference should therefore not be interpreted as evidence of equivalence. More generally, the small infection and hybrid-exposure groups limited precision and the ability to account comprehensively for potentially important covariates. The intensive follow-up required for this birth cohort may also have preferentially retained highly adherent families, potentially limiting generalizability beyond similarly engaged US populations.

The present findings also do not establish the clinical consequences of the observed antibody trajectories. The study was not designed to measure vaccine effectiveness or protection from infection or disease, and antibody magnitude or persistence cannot be interpreted as a direct measure of clinical protection. Likewise, the absence of direct cellular measurements means that mechanisms proposed to explain exposure-associated differences, including B-cell imprinting, affinity maturation, or long-lived plasma-cell formation, cannot be inferred from these data. Decay estimates were dependent on the number and timing of longitudinal samples, and estimates with wide confidence intervals require particular caution. The absence of measurable decay should not be interpreted as indefinite persistence.

Taken together, these findings indicate that the nature and sequence of influenza exposure during early life are associated with distinct longitudinal serum antibody trajectories. Primary infection was associated with greater-magnitude responses and less measurable decline than primary vaccination in this cohort, whereas repeated vaccination was associated with progressively greater antibody responses. Among children with prior infection, subsequent vaccination was associated with an initial bias toward the previously encountered subtype among the HA antigens tested, which became less pronounced with repeated vaccination. These observations provide longitudinal evidence that early-life exposure history is associated with the subsequent magnitude, persistence, and antigenic distribution of serum antibody responses. Further studies incorporating cellular measurements, antigenically matched infecting and assay viruses, and clinical outcomes will be important for determining the mechanisms and protective relevance of these exposure-associated antibody trajectories.

## METHODS

### Study cohort

The longitudinal IMPRINT influenza cohort aims to understand how initial and subsequent influenza infections and/or vaccinations shape early life immunity. The cohort includes mother-child pairs followed from the third trimester through the first years of childhood, with regularly collected blood samples, weekly nasal swabs, twice-weekly symptom surveys, and clinical/epidemiologic data collection to detect and characterize respiratory infections in early life. IMPRINT received Institutional Review Board approval at the Cincinnati Children’s Hospital Medical Center (Cincinnati, OH) (IRB 2019-0629) and the University of Cincinnati Medical Center (Cincinnati, OH) (IRB SITE00000489); written informed consent for the mother and child was obtained from each participating mother at their enrollment visit.

All the experiments were performed in plasma or serum samples collected longitudinally from participants and cryopreserved. The peripheral blood samples used in this analysis were collected longitudinally starting at birth, at week 6 of life, each summer, and after reported influenza exposures (vaccination and/or infection). Our sample of cohort participants included 45 males and 46 females, having high adherence to study protocols and specimen/data collection. The description of study participants is provided in **Table 1**.

### Cell lines

The Madin-Darby Canine Kidney Cells (MDCK) and MDCK expressing Sialyltransferase-1 (MDCK-SIAT)^37^ cells were a generous gift from Dr. Anice Lowen and were cultured in complete Dulbecco’s Modified Eagle Medium (DMEM) supplemented with 10% fetal bovine serum, and 100U/ml penicillin/streptomycin^38–40^.

### Viruses

The influenza viruses A/Wisconsin/2019 (H1N1), A/Wisconsin/2022 (H1N1), and A/Darwin/2021 (H3N2) were handled under BSL2 conditions. The H1N1 viruses were propagated in 9-11-day-old embryonated chicken eggs and incubated for 48 h at 37 °C. Allantoic fluid was collected and used as the viral stock for experiments. H3N2 virus was propagated in MDCK-SIAT cells containing 1 ug/mL TPCK-treated Trypsin and stored at −80 C until use.

### ELISA

ELISAs were performed as described previously^41^ with a few modifications. Briefly, influenza HA antigen at 1 ug/mL in 1X PBS was coated on MaxiSorp plates (Thermo Fisher Scientific, #439454) at 4°C overnight. The plates were blocked with 1% BSA and 0.05% Tween-20 in 1X PBS for 2 h. Three-fold serially diluted plasma was added to the plates and incubated at room temperature for 1 hour. After incubation, the plates were washed 3 times with PBS containing 0.05% Tween 20, and influenza vaccine-specific IgG signals were detected by incubation with horseradish peroxidase (HRP)–conjugated anti-human IgG (Jackson ImmunoResearch Laboratories, #109-036-098). Plates were then washed thoroughly and developed with o-phenylenediamine substrate (Sigma-Aldrich, #P8787) in 0.05 M phosphate-citrate buffer (Sigma-Aldrich, #P4809), pH 5.0, containing 0.012% hydrogen peroxide (Thermo Fisher Scientific, #18755). Absorbance was measured at 490 nm.

### HAI assay

The HAI assays were performed as described^42,43^ in 96-well V-bottom microtiter plates (VWR #76446-958). To remove nonspecific inhibitors of HA, sera were incubated at 37°C overnight (18 ± 1 hr) at a 1:4 dilution with receptor-destroying enzyme (RDE; Denka Seiken, Tokyo, Japan), followed by a 45-min inactivation in a 56°C water bath and further dilution to 1:10 with 0.9 % saline. The virus inoculum was back-titrated right before the HAI assay to confirm the precision of the HA units. 25 uL of 2-fold serially diluted plasma (starting from 1:10) was incubated with 25 ul of 4 HA units of virus in V-bottom microtiter plates for 30 minutes at RT. A 50-ul volume of 0.5% turkey RBCs was added, and the reaction mixture was incubated for 1 hr at RT. Wells were examined visually for inhibition of HA, as indicated by the appearance of well-defined RBC “buttons” or streak formation upon plate tilting. HAI titers were defined as the reciprocal of the highest serum dilution that completely prevented HA.

### Focus Reduction Neutralization (FRNT) Assay

The FRNT assays were performed as described previously^44,45^. Briefly, the plasma samples were incubated with receptor-destroying enzyme (RDE; Denka Seiken, Tokyo, Japan) at a 1:4 dilution overnight (18 ± 1 hr) at 37 °C, followed by a 45-min inactivation in a 56 °C water bath and further dilution to 1:10 with 0.9% saline. The plasma samples were serially diluted 2-fold in serum-free DMEM containing 1 ug/mL TPCK-treated trypsin (Sigma #T1426), starting with an initial dilution of 1:10 in a total volume of 60 uL, followed by incubation with an equal volume of the respective influenza virus. The plasma-virus mixture was incubated for 1 hr at 37 ᵒC and then added to MDCK cells seeded the previous day at a density of 2.5 × 10^4^ cells/well in a 96-well cell culture plate. After 1 hr of incubation, the antibody-virus mixture was removed, and a pre-warmed overlay of 0.8% methylcellulose, 1% BSA, and 1 ug/mL TPCK-treated trypsin in DMEM was added to each well. The cells were incubated for 8-30 hr at 37 ᵒC and washed 4 times with 1X PBS. The cells were fixed with a 1:1 mixture of acetone and methanol for 30 min. Following fixation, the cells were washed and blocked with 5 % non-fat dry milk in 1X PBS. 1 ug/mL of CR9114 IgG1 antibody^46^ in 5 % non-fat dry milk was added and incubated at RT for 2 hr. CR9114 IgG1 was used as the detection antibody to identify influenza HA-expressing infected cells/foci. The plates were washed 3x with 1X PBS, and horseradish peroxidase (HRP)–conjugated anti-human IgG diluted to 1:10000 in 5 % non-fat dry milk was added. Following 1 hr incubation, the plates were washed 3x with 1X PBS, developed with TrueBlue substrate, and read on an ELISpot reader. The FRNT titers for 50% virus inhibition are reported. Percent neutralization was calculated as 1 − (mean foci in serum-containing wells / mean foci in virus-control wells) × 100. FRNT50 was defined as the reciprocal serum dilution associated with 50% reduction in focus number relative to the virus-control condition and was determined from the dilution-response curve.

### Quantification and statistical analysis

The data were analyzed using GraphPad Prism 10.6.1. The antibody neutralization titers were quantified by counting foci per sample. The neutralization titers were calculated as [1 – (average number of foci in wells incubated with patient serum) ÷ (average number of foci in wells incubated with control serum)].

Antibody decay curves were modeled using a mixed-effects exponential decay model implemented in MonolixSuite 2024R1. For each specified condition, fitting was performed only using measurements obtained at or after day 28 post-infection or post-vaccination from individuals with more than two such measurements.

## Acknowledgements

We thank the participants and parents of infant donors for volunteering their time and effort to take part in this study. This work was supported in part by grants U01AI144673-02, 3U19AI057266-17S2, and U54CA260563 from the National Institute of Allergy and Infectious Diseases (NIAID), National Institutes of Health (NIH), and by the Oliver S. and Jennie R. Donaldson Charitable Trust, Emory Executive Vice President for Health Affairs Synergy Fund award, the Georgia Research Alliance, the Pediatric Research Alliance Center for Childhood Infections and Vaccines and Children’s Healthcare of Atlanta, and a Woodruff Health Sciences Center 2020 COVID-19 CURE Award.

## Author contributions

D.J. performed the design of the study, data acquisition, data curation and analysis, data visualization, and wrote the original draft of the manuscript. S.K. and V.Z. contributed to data curation and analysis, data visualization, writing review, and editing. M.A.S., A.B., and C.R. created the IMPRINT cohort, recruited and enrolled mothers and their children, conducted study visits, collected clinical data and samples for the study, and curated data for the analyses conducted. S.B. assisted in virus growth and amplification. T.C., F.A.M., and L.X. contributed to data curation and review of the manuscript. A.L., M.A.S., and J.W. contributed to the conception and design of the study and the obtaining of study funding. All authors contributed to editing the manuscript. All authors have read and accepted the manuscript.

## Declaration of interests

All authors declare no conflicts of interest.

---

**Supplementary Figure 1:**
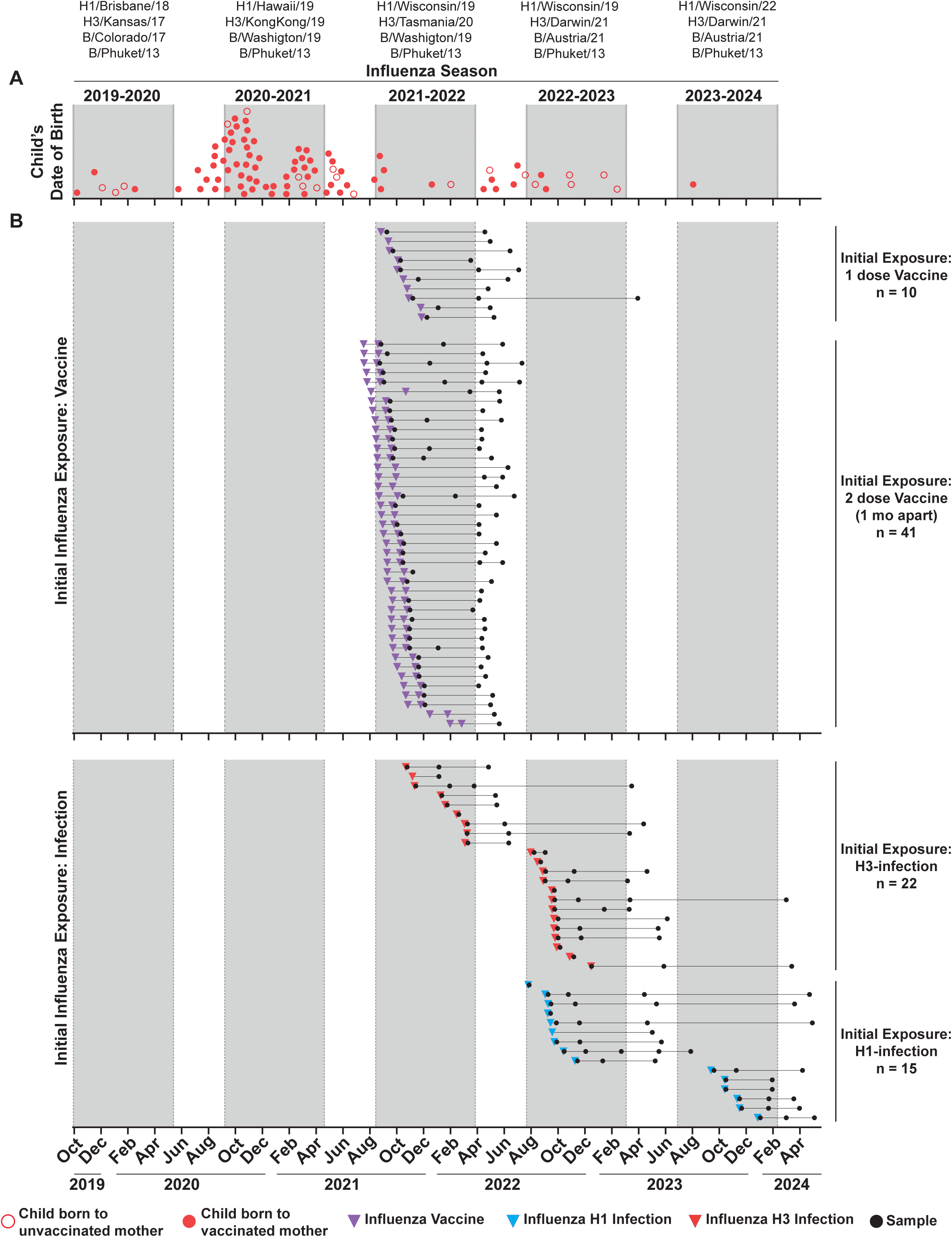
Layout of infants included in the study along with their initial influenza exposure history and samples included in the analysis. (A) Infants included in the in the study along with their date of birth and maternal vaccination status. (B) Infants receiving influenza-vaccination as an intial influenza exposure along with the timing of vaccination and timing of longitudinal sample collections for each infant. (C) Infants with influenza-infection as an intial influenza exposure along with the timing of H1- or H3-infection and timing of longitudinal sample collections for each infant.

**Supplementary Figure 2:**
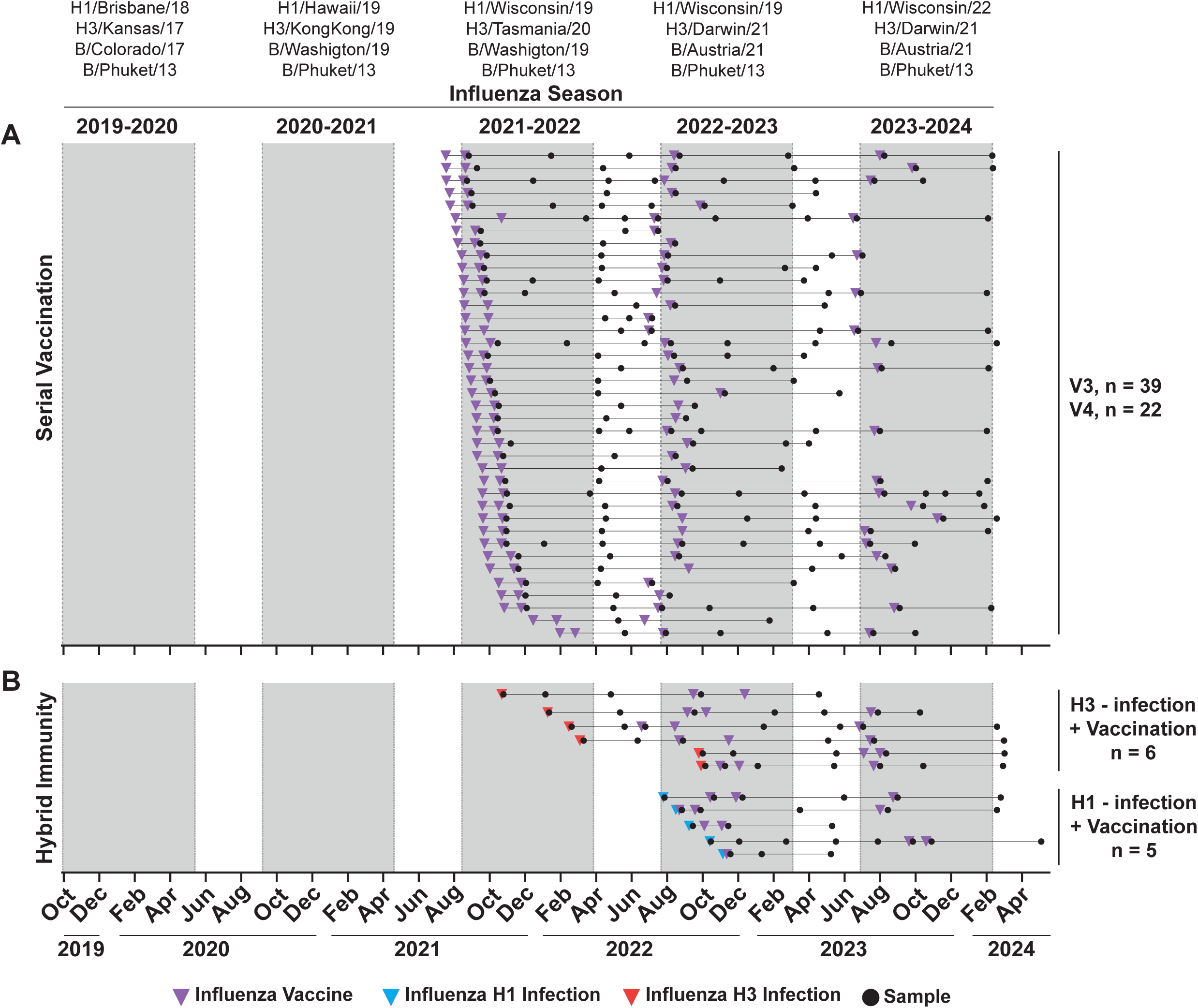
Layout of infants included to study multiple influenza exposures along with their exposure history and samples included in the analysis. (A) Infants included to study serial influenza vaccination along with the vaccination history and timing of longitudinal sample collections for each infant. (C) Infants included to study hybrid immune responses as influenza-infection followed by vaccination along with their influenza exposure histort and timing of longitudinal sample collections for each infant.

## REFERENCES

1. Ortiz, J.R., Englund, J.A., and Neuzil, K.M. (2011). Influenza vaccine for pregnant women in resource-constrained countries: a review of the evidence to inform policy decisions. Vaccine 29, 4439–4452.

2. Boudreau, C.M., Burke, J.S., Shuey, K.D., Wolf, C., Katz, J., Tielsch, J., Khatry, S., LeClerq, S.C., Englund, J.A., and Chu, H.Y. (2022). Dissecting Fc signatures of protection in neonates following maternal influenza vaccination in a placebo-controlled trial. Cell Reports 38.

3. Ranjeva, S., Subramanian, R., Fang, V.J., Leung, G.M., Ip, D.K., Perera, R.A., Peiris, J.M., Cowling, B.J., and Cobey, S. (2019). Age-specific differences in the dynamics of protective immunity to influenza. Nature communications 10, 1660.

4. Francis, T. (1960). On the doctrine of original antigenic sin. Proceedings of the American Philosophical Society 104, 572–578.

5. Zhang, A., Stacey, H.D., Mullarkey, C.E., and Miller, M.S. (2019). Original antigenic sin: how first exposure shapes lifelong anti–influenza virus immune responses. The Journal of Immunology 202, 335–340.

6. Tesini, B.L., Kanagaiah, P., Wang, J., Hahn, M., Halliley, J.L., Chaves, F.A., Nguyen, P.Q., Nogales, A., DeDiego, M.L., and Anderson, C.S. (2019). Broad hemagglutinin-specific memory B cell expansion by seasonal influenza virus infection reflects early-life imprinting and adaptation to the infecting virus. Journal of virology 93, 10.1128/jvi.00169-00119.

7. Gostic, K.M., Bridge, R., Brady, S., Viboud, C., Worobey, M., and Lloyd-Smith, J.O. (2019). Childhood immune imprinting to influenza A shapes birth year-specific risk during seasonal H1N1 and H3N2 epidemics. PLoS pathogens 15, e1008109.

8. Knight, M., Changrob, S., Li, L., and Wilson, P.C. (2020). Imprinting, immunodominance, and other impediments to generating broad influenza immunity. Immunological reviews 296, 191–204.

9. Edler, P., Schwab, L.S., Aban, M., Wille, M., Spirason, N., Deng, Y.-M., Carlock, M.A., Ross, T.M., Juno, J.A., and Rockman, S. (2024). Immune imprinting in early life shapes cross-reactivity to influenza B virus haemagglutinin. Nature Microbiology 9, 2073–2083.

10. Nelson, S.A., and Sant, A.J. (2019). Imprinting and editing of the human CD4 T cell response to influenza virus. Frontiers in Immunology 10, 932.

11. Sherman, A.C., Lai, L., Bower, M., Natrajan, M.S., Huerta, C., Karmali, V., Kleinhenz, J., Xu, Y., Rouphael, N., and Mulligan, M.J. (2020). The effects of imprinting and repeated seasonal influenza vaccination on adaptive immunity after influenza vaccination. Vaccines 8, 663.

12. https://www.cdc.gov/covid/index.html.

13. (U.S.), C.f.D.C.a.P. Research use only CDC influenza SARS-CoV-2 (Flu SC2) multiplex assay real-Time RT-PCR primers and probes.

14. Gonsalves, S., Mahony, J., Rao, A., Dunbar, S., and Juretschko, S. (2019). Multiplexed detection and identification of respiratory pathogens using the NxTAG® respiratory pathogen panel. Methods 158, 61–68. 10.1016/j.ymeth.2019.01.005.

15. Zhong, Z., Haltalli, M., Holder, B., Rice, T., Donaldson, B., O’driscoll, M., Le-Doare, K., Kampmann, B., and Tregoning, J. (2019). The impact of timing of maternal influenza immunization on infant antibody levels at birth. Clinical & Experimental Immunology 195, 139–152.

16. Li, M., Wang, W., Chen, J., Zhan, Z., Xu, M., Liu, N., Ren, L., You, L., Zheng, W., and Shi, H. (2023). Transplacental transfer efficiency of maternal antibodies against influenza A (H1N1) pdm09 virus and dynamics of naturally acquired antibodies in Chinese children: a longitudinal, paired mother–neonate cohort study. The Lancet Microbe 4, e893–e902.

17. Kelvin, A.A., and Zambon, M. (2019). Influenza imprinting in childhood and the influence on vaccine response later in life. Eurosurveillance 24, 1900720.

18. Miller, M.S., Gardner, T.J., Krammer, F., Aguado, L.C., Tortorella, D., Basler, C.F., and Palese, P. (2013). Neutralizing antibodies against previously encountered influenza virus strains increase over time: a longitudinal analysis. Science translational medicine 5, 198ra107–198ra107.

19. Daulagala, P., Mann, B.R., Leung, K., Lau, E.H., Yung, L., Lei, R., Nizami, S.I., Wu, J.T., Chiu, S.S., and Daniels, R.S. (2023). Imprinted anti-hemagglutinin and anti-neuraminidase antibody responses after childhood infections of A (H1N1) and A (H1N1) pdm09 influenza viruses. MBio 14, e00084–00023.

20. Brouwer, A.F., Balmaseda, A., Gresh, L., Patel, M., Ojeda, S., Schiller, A.J., Lopez, R., Webby, R.J., Nelson, M.I., and Kuan, G. (2022). Birth cohort relative to an influenza A virus’s antigenic cluster introduction drives patterns of children’s antibody titers. PLoS Pathogens 18, e1010317.

21. Maltseva, M., Keeshan, A., Cooper, C., and Langlois, M.-A. (2024). Immune imprinting: The persisting influence of the first antigenic encounter with rapidly evolving viruses. Human vaccines & immunotherapeutics 20, 2384192.

22. Joshi, D., Nyhoff, L.E., Zarnitsyna, V.I., Moreno, A., Manning, K., Linderman, S., Burrell, A.R., Stephens, K., Norwood, C., and Mantus, G. (2023). Infants and young children generate more durable antibody responses to SARS-CoV-2 infection than adults. Iscience 26.

23. Wimmers, F., Burrell, A.R., Feng, Y., Zheng, H., Arunachalam, P.S., Hu, M., Spranger, S., Nyhoff, L.E., Joshi, D., and Trisal, M. (2023). Multi-omics analysis of mucosal and systemic immunity to SARS-CoV-2 after birth. Cell 186, 4632–4651. e4623.

24. Kuraoka, M., Curtis, N.C., Watanabe, A., Tanno, H., Shin, S., Ye, K., Macdonald, E., Lavidor, O., Kong, S., and Von Holle, T. (2022). Infant antibody repertoires during the first two years of influenza vaccination. Mbio 13, e02546–02522.

25. Worobey, M., Plotkin, S., and Hensley, S.E. (2020). Influenza vaccines delivered in early childhood could turn antigenic sin into antigenic blessings. Cold Spring Harbor perspectives in medicine 10, a038471.

26. Koutsakos, M., Nguyen, T.H., and Kedzierska, K. (2019). With a little help from T follicular helper friends: humoral immunity to influenza vaccination. The Journal of Immunology 202, 360–367.

27. Sangster, M.Y., Nguyen, P.Q., and Topham, D.J. (2019). Role of memory B cells in hemagglutinin-specific antibody production following human influenza A virus infection. Pathogens 8, 167.

28. Kumagai, T., Nagai, K., Okui, T., Tsutsumi, H., Nagata, N., Yano, S., Nakayama, T., Okuno, Y., and Kamiya, H. (2004). Poor immune responses to influenza vaccination in infants. Vaccine 22, 3404–3410.

29. Alexander-Miller, M.A. (2020). Challenges for the newborn following influenza virus infection and prospects for an effective vaccine. Frontiers in immunology 11, 568651.

30. Chiu, C., Ellebedy, A.H., Wrammert, J., and Ahmed, R. (2014). B cell responses to influenza infection and vaccination. Influenza Pathogenesis and Control-Volume II, 381–398.

31. https://www.cdc.gov/vaccines/hcp/imz-schedules/child-adolescent-notes.html. (2025).

32. Davis, C.W., Jackson, K.J., McCausland, M.M., Darce, J., Chang, C., Linderman, S.L., Chennareddy, C., Gerkin, R., Brown, S.J., and Wrammert, J. (2020). Influenza vaccine–induced human bone marrow plasma cells decline within a year after vaccination. Science 370, 237–241.

33. Clemens, E.A., and Alexander-Miller, M.A. (2021). Understanding antibody responses in early life: baby steps towards developing an effective influenza vaccine. Viruses 13, 1392.

34. Crofts, K.F., Holbrook, B.C., Page, C.L., Gillespie, R.A., D’Agostino Jr, R.B., Sangesland, M., Ornelles, D.A., Kanekiyo, M., and Alexander-Miller, M.A. (2025). Antibody function predicts viral control in newborn monkeys immunised with an influenza virus HA stem nanoparticle. Nature Communications 16, 3785.

35. Nunes, M.C., Cutland, C.L., Jones, S., Hugo, A., Madimabe, R., Simões, E.A., Weinberg, A., and Madhi, S.A. (2016). Duration of infant protection against influenza illness conferred by maternal immunization: secondary analysis of a randomized clinical trial. JAMA pediatrics 170, 840–847.

36. Sugimura, T., Ito, Y., Tananari, Y., Ozaki, Y., Maeno, Y., Yamaoka, T., and Kudo, Y. (2008). Improved antibody responses in infants less than 1 year old using intradermal influenza vaccination. Vaccine 26, 2700–2705.

37. Matrosovich, M., Matrosovich, T., Carr, J., Roberts, N.A., and Klenk, H.-D. (2003). Overexpression of the α-2, 6-sialyltransferase in MDCK cells increases influenza virus sensitivity to neuraminidase inhibitors. Journal of virology 77, 8418–8425.

38. Powell, H., and Pekosz, A. (2020). Neuraminidase antigenic drift of H3N2 clade 3c. 2a viruses alters virus replication, enzymatic activity and inhibitory antibody binding. PLoS pathogens 16, e1008411.

39. Lee, L.Y., Zhou, J., Koszalka, P., Frise, R., Farrukee, R., Baba, K., Miah, S., Shishido, T., Galiano, M., and Hashimoto, T. (2021). Evaluating the fitness of PA/I38T-substituted influenza A viruses with reduced baloxavir susceptibility in a competitive mixtures ferret model. PLoS pathogens 17, e1009527.

40. Yegorov, S., Celeste, D.B., Gomes, K.B., Ang, J.C., Vandenhof, C., Wang, J., Rybkina, K., Tsui, V., Stacey, H.D., and Loeb, M. (2022). Inactivated and live-attenuated seasonal influenza vaccines boost broadly neutralizing antibodies in children. Cell Reports Medicine 3.

41. Kumar, S., Patel, A., Lai, L., Chakravarthy, C., Valanparambil, R., Reddy, E.S., Gottimukkala, K., Davis-Gardner, M.E., Edara, V.V., and Linderman, S. (2022). Structural insights for neutralization of Omicron variants BA. 1, BA. 2, BA. 4, and BA. 5 by a broadly neutralizing SARS-CoV-2 antibody. Science advances 8, eadd2032.

42. Lowen, A.C., Steel, J., Mubareka, S., Carnero, E., García-Sastre, A., and Palese, P. (2009). Blocking interhost transmission of influenza virus by vaccination in the guinea pig model. Journal of virology 83, 2803–2818.

43. Zacour, M., Ward, B.J., Brewer, A., Tang, P., Boivin, G., Li, Y., Warhuus, M., McNeil, S.A., LeBlanc, J.J., and Hatchette, T.F. (2016). Standardization of hemagglutination inhibition assay for influenza serology allows for high reproducibility between laboratories. Clinical and Vaccine Immunology 23, 236–242.

44. Gouma, S., Kim, K., Weirick, M.E., Gumina, M.E., Branche, A., Topham, D.J., Martin, E.T., Monto, A.S., Cobey, S., and Hensley, S.E. (2020). Middle-aged individuals may be in a perpetual state of H3N2 influenza virus susceptibility. Nature communications 11, 4566.

45. Furey, C., Scher, G., Ye, N., Kercher, L., DeBeauchamp, J., Crumpton, J.C., Jeevan, T., Patton, C., Franks, J., and Rubrum, A. (2024). Development of a nucleoside-modified mRNA vaccine against clade 2.3. 4.4 b H5 highly pathogenic avian influenza virus. Nature Communications 15, 4350.

46. Guthmiller, J.J., Han, J., Utset, H.A., Li, L., Lan, L.Y.-L., Henry, C., Stamper, C.T., McMahon, M., O’Dell, G., Fernández-Quintero, M.L., et al. (2022). Broadly neutralizing antibodies target a haemagglutinin anchor epitope. Nature 602, 314–320. 10.1038/s41586-021-04356-8.

